# CASPR facilitates clearance of degenerating axons by Schwann cells and macrophages after peripheral nerve injury

**DOI:** 10.1101/2025.10.19.683281

**Authors:** Sofia Meyer zu Reckendorf, Anna-Luisa Klein, Antonia Seitz, Jutta Hegler, Claudia Klugmann, Daniela Sinske, Steffanie Deininger, Maria Pedro, Bernd Knöll

## Abstract

In the peripheral nervous system (PNS) debris of injured axons is efficiently removed by Schwann cells (SCs) and macrophages (Mphs) – a process which is vital for nerve regeneration. Molecular so-called “eat-me” signals, mediating axon-SC/axon-Mph crosstalk and debris clearance after injury are just at the beginning of being identified. Herein, we describe the axonal node protein contactin-associated protein (CASPR) as a potential novel signal for axonal debris removal in peripheral nerve injury. In healthy nerves, CASPR is restricted to Ranvier’s nodes interacting with glia-derived partner proteins to assure saltatory action potential propagation. In injured murine and human nerves, we describe upregulation and re-localization of CASPR protein along the axons. Enhanced CASPR presence after injury appears to involve local axonal translation rather than transcriptional regulation. Of note, CASPR overexpression in murine central and peripheral neurons *in vitro* has a growth inhibitory effect. More importantly, axonal debris deriving from injured nerves – and hence having increased CASPR content – are phagocytosed more efficiently than debris of healthy nerves. Interfering with CASPR by function-blocking antibodies strongly reduces axonal debris uptake. This finding demonstrates a functional relevance of CASPR as a potential “eat me” signal in this process.

## Introduction

In the peripheral nervous system (PNS) injured axons have a remarkable potential for regeneration, which is not matched by axons in the central nervous system (CNS). In injured peripheral nerves, Schwann cells (SCs), along with other cells (e.g. macrophages; Mphs), create a growth-permissive environment for injured axons (1). After nerve injury, SCs undergo a phenotypic conversion to repair SCs, which promote axonal regeneration and target reinnervation. Repair SCs provide growth factors (e.g. GDNF, NGF, BDNF) and form Büngner bands as spatial cues to support and guide axonal regrowth (1, 2). Importantly, repair SCs also contribute to myelin and axon debris removal and attract Mphs, which – as professional phagocytes – accelerate debris clearance (1–4). In fact, clearance of debris after injury seems to be a crucial process preceding regeneration, since blocking the trans-differentiation of SCs into repair SCs and hence blocking inflammation and debris removal, hinders regeneration (2, 3).

The precise orchestration of inflammation, debris clearance and axonal outgrowth is crucial for nerve regeneration, yet the complex interactions between phagocytosing cells (Schwann cells and macrophages) and neurons/axons during debris uptake remain poorly understood. Outside the nervous system, it is well established that apoptotic cells present “eat-me” signals on their surface to trigger phagocytosis (5–7). One of the best-characterized “eat-me” signals is phosphatidylserine (PS), which engages phagocytes via PS receptors, TAM receptors (Tyro-3, AXL, MerTK) via bridging proteins Gas6 or protein S, or through integrin receptors via MFG-E8. Additionally, membrane glycoproteins bearing mannose, fucose, or N-acetylglucosamine can activate the mannose receptor (MR/CD206) to promote uptake (7, 8). In peripheral nerve injury, although the precise mechanisms remain unclear, degenerating axons have been shown to expose PS, while Schwann cells express PS receptors, TAM receptors, and the MR (7–10).

This study investigates the role of CASPR during the acute injury response in peripheral nerves. CASPR is well known as an axonal structural component anchoring the myelin sheath to the axon at paranodal regions of the nodes of Ranvier (11–13). Specifically, axonal CASPR interacts with Neurofascin (NFASC) 155 expressed on SCs through axonal Contactin (CNTN) proteins. Here, we describe that axons undergoing Wallerian degeneration after injury upregulate CASPR through local translation and redistribute CASPR along the axon. Notably, this mechanism is highly conserved between mouse and human nerves. Further, we show that this local CASPR upregulation most likely suppresses axonal outgrowth in this acute injury phase. Finally, we describe a novel CASPR-mediated axon-SC and axon-Mph cross-talk mechanism that promotes axonal debris uptake.

Together, these findings identify CASPR as a novel axonal “eat-me” signal, which is induced upon nerve injury to facilitate efficient phagocytosis of axonal debris by SCs and Mphs.

## Results

### Nerve injury results in CASPR upregulation and re-distribution in murine and human nerves

To study changes in CASPR expression and localization after nerve injury, an *in vivo* sciatic nerve crush model in mice was employed and CASPR expression was quantified 3 days after injury (Fig. 1). In longitudinal sections of uninjured nerves, CASPR was expectedly restricted to nodes of Ranvier (Fig. 1A, A’). In contrast, 3 days after injury, CASPR expression was increased and spread laterally along axons (Fig. 1B-B’’’). CASPR upregulation and redistribution was visible in the distal stump and in the proximal stump close to the injury site, which are the regions undergoing Wallerian degeneration (WD) after injury (Fig. 1B-B’’’). In contrast, proximal axon regions more distant to the injury site – hence not undergoing WD – did not show altered CASPR abundance or localization (Fig. 1B’).

**Figure 1:**
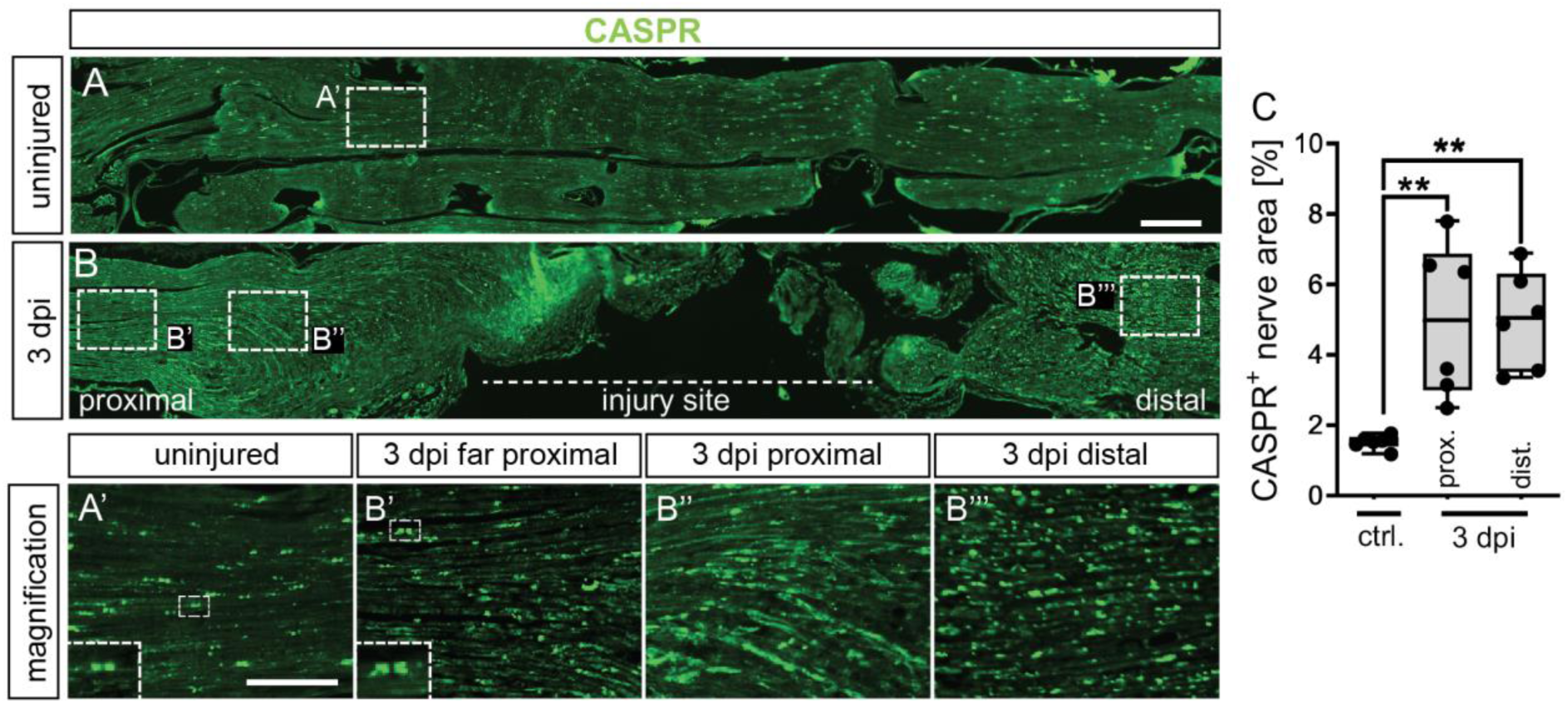
Peripheral nerve injury in mice induces CASPR upregulation in degenerating axons. (A-B) CASPR staining in longitudinal sections of murine sciatic nerves without injury (A) or at 3 days post injury (B). (A’, B’-B’’’) show higher magnifications of the areas indicated in (A) and (B). (C) Quantification of CASPR^+^ nerve area in uninjured vs. injured nerves at proximal (corresponding to B’’) and distal (corresponding to B’’’) regions. Each dot in the graph represents one animal (n = 6). Data are presented as box plots including the mean and whiskers min to max. Statistical analysis was performed using a two-sided Mann-Whitney test with *p < 0.05; **p < 0.01; ***p < 0.001. Scale bar in (A) applies for (A-B) and corresponds to 100 µm. Scale bar in (A’) applies for (A’-B’’’) and corresponds to 40 µm. CASPR: Contactin associated protein 1, dpi: days post injury

The observed CASPR upregulation in regions of WD was somewhat unexpected, since axons in those regions degenerate and myelin sheaths are degraded, making CASPR presence as structural myelin anchoring protein obsolete. In order to clarify if those changes were specific to the sciatic nerve or rather a general event in the PNS, we compared CASPR abundance between injured sciatic and facial nerves. Here, cross sections of healthy and injured facial nerves likewise presented CASPR upregulation (Fig. S1), pointing at a conserved mechanism between different PNS nerves.

Next, we utilized our previously reported *ex vivo* injury model (14) to determine whether the *in vivo* CASPR upregulation findings (Fig. 1) can be replicated *ex vivo* (Fig. 2). Freshly dissected sciatic nerves were cultured for up to 48 hours to monitor acute injury responses (16). This system is also applicable to human nerves, enabling a direct comparison of human and murine injury responses (16), which we included in this study (Fig. 2).

**Figure 2:**
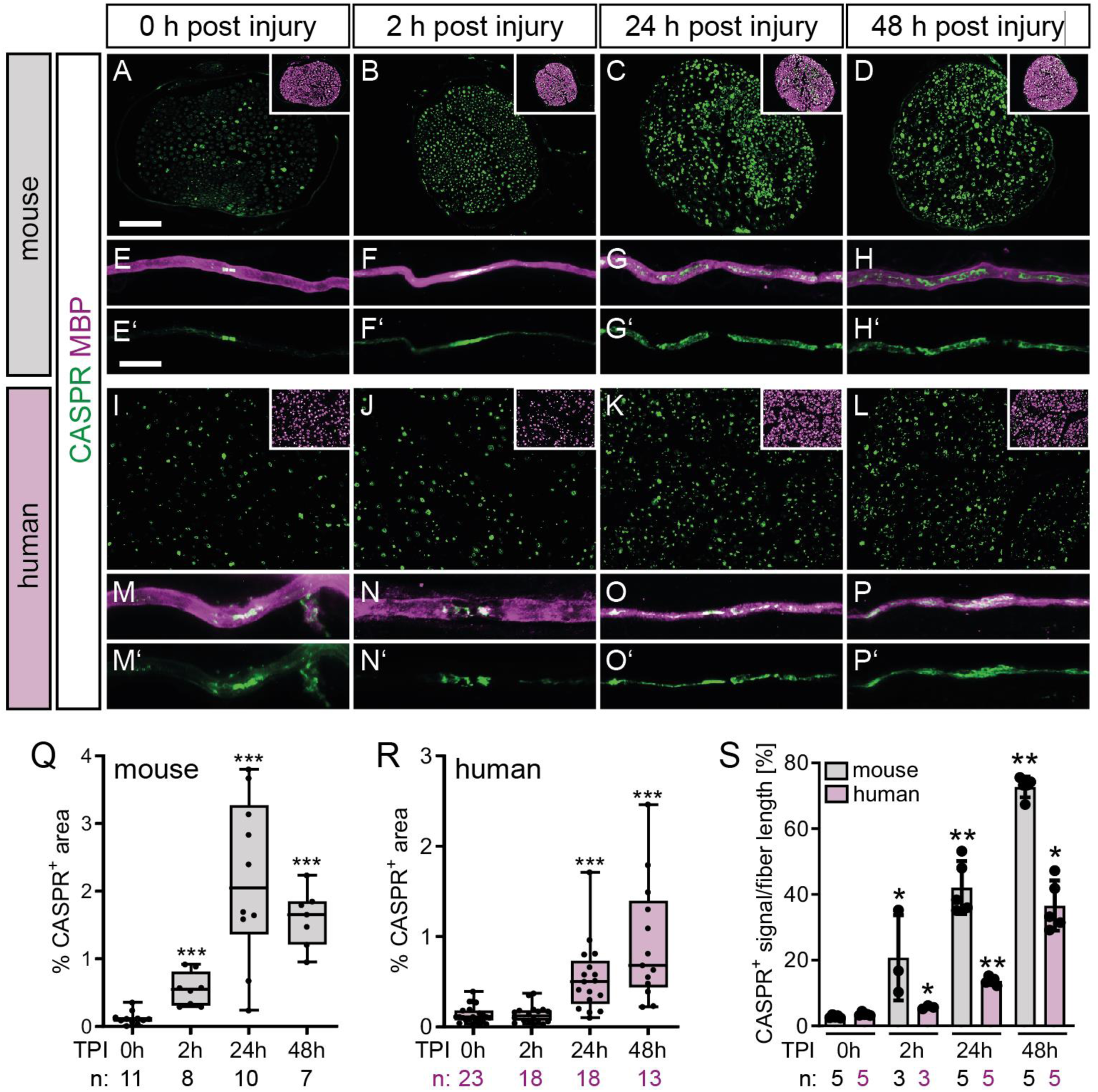
Injured murine and human nerve explants show CASPR upregulation and redistribution along degenerating axons. (A-D) and (I-L) Cross sections of murine (A-D) and human (I-L) sural nerve explants without injury (0 h post injury) and at different time points after injury as indicated. Pictures show CASPR staining while inserts depict the overlay of CASPR with MBP. (E-H) and (M-P) Teased fibers of murine (E-H) and human (M-P) sural nerves without injury (0 h post injury) and at different time points after injury as indicated. Staining depicts the overlay of CASPR and MBP. Separate CASPR channel is depicted in (E’-H’) and (M’-P’). (Q-R) Quantification of CASPR stained area in murine (Q) and human (R) sections. (S) Quantification of CASPR positive area normalized to fiber length in teased fibers of murine and human sural nerves. Each dot in the graphs represents one animal or patient sample. Numbers of samples (n) are indicated below each graph. Data in (Q-R) are presented as box plots including the mean and whiskers min to max. Bars in (S) represent mean values with SD. Statistical analysis was performed using a two-sided Mann–Whitney test with *p < 0.05; **p < 0.01; ***p < 0.001. Scale bar in (A) applies for (A-D) and (I-L) and corresponds to 50 µm. Scale bar in (E) applies for (E-H) and (M-P) and corresponds to 25 µm. CASPR: contactin associated protein 1, MBP: myelin basic protein, TPI: time post injury

First, we investigated CASPR expression in murine sciatic nerves 2 h, 24 h and 48 h after dissection/injury (Fig. 2A-D, Q, S). We used nerve cross-sections (Fig. 2A-D) and additionally single teased fibers to provide a longitudinal view on individual nodes (Fig. 2E-H). As observed *in vivo* (Fig. 1), CASPR was also induced in this *ex vivo* injury model, starting already at 2 h after injury (Fig. 2A-B, Q). This was even more pronounced at 24 h and 48 h post injury (Fig. 2C-D, Q). Furthermore, teased fibers confirmed the lateral spread of CASPR along the axons in contrast to the un-injured condition (Fig. 2E-H’).

Next, this injury-induced CASPR upregulation in mice was also analyzed in human sural nerves (Fig. 2I-L, M-P; R, S). Therefore, human sural nerves, collected from patients during reconstructive surgery were employed as before (19; Table 1). In un-injured nerves, CASPR localization was restricted to the nodes (Fig. 2M). As observed in rodents, CASPR was also upregulated in human nerves upon injury (Fig. 2I-L, R). However, this response was delayed in humans compared to mice, as it took 24h for the upregulation to become visible compared to 2h in mice (Fig. 2S).

**Table 1:** Patient information.

| *Pat. ID* | *Age* | *Sex* | *Sample used for* |
| --- | --- | --- | --- |
| P1 | 61 | f | HIS |
| P2 | 53 | m | HIS |
| P3 | 72 | f | HIS |
| P4 | 50 | m | HIS |
| P5 | 66 | f | HIS |
| P6 | 57 | f | HIS |
| P7 | 36 | m | HIS |
| P8 | 54 | m | HIS |
| P9 | 62 | m | HIS |
| P10 | 51 | m | HIS |
| P11 | 62 | f | HIS |
| P12 | 66 | f | HIS |
| P13 | 65 | f | HIS |
| P14 | 20 | f | HIS, WB |
| P15 | 26 | m | WB |
| P16 | 61 | m | HIS |
| P17 | 51 | m | HIS, WB |
| P18 | 67 | m | HIS |
| P19 | 72 | m | HIS |
| P20 | 41 | m | HIS |
| P21 | 63 | m | HIS |
| P22 | 61 | m | HIS, WB |
| P23 | 61 | f | HIS, TF |
| P24 | 45 | m | HIS, TF |
| P25 | 57 | m | HIS |
| P26 | 53 | f | TF, TF It, WB It, |
| P27 | 47 | m | TF, TF It, WB It, |
| P28 | 66 | m | WB It |
| P29 | 23 | m | TF It |
| P30 | 41 | m | WB It |
| P31 | 18 | m | TF It |
| P32 | 36 | m | TF |
| P33 | 54 | f | WB |
| P34 | 55 | m | WB |
| P35 | 65 | m | WB |
| P36 | 49 | f | WB |
| P37 | 59 | f | WB |
| P38 | 23 | f | SC |
| P39 | 75 | f | SC |

Abbreviations: HIS, histology; SC, Schwann cell cultures; TF, teased fibers; TF It, Teased fibers local translation; WB, western blot; IB It, immunoblot local translation

In summary, *in vivo* and *ex vivo* analysis of injured rodent and human peripheral nerves revealed CASPR upregulation and lateral spreading along the injured axons undergoing WD.

### CASPR expression is increased in injured murine and human peripheral nerves together with CNTN1 and NFASC 155

Above we described CASPR upregulation in murine and human peripheral nerves upon injury (Fig. 1 and 2). Since this histological analysis cannot unequivocally distinguish between CASPR upregulation and/or a mere re-distribution we used nerve explants, prepared protein lysates and performed western blot analysis (Fig. 3). In both, murine and human nerves, CASPR abundance was elevated after injury compared to uninjured nerves, although human nerves again showed slightly delayed injury responses compared to mice (48 h vs. 24 h; Fig. 3A).

**Figure 3:**
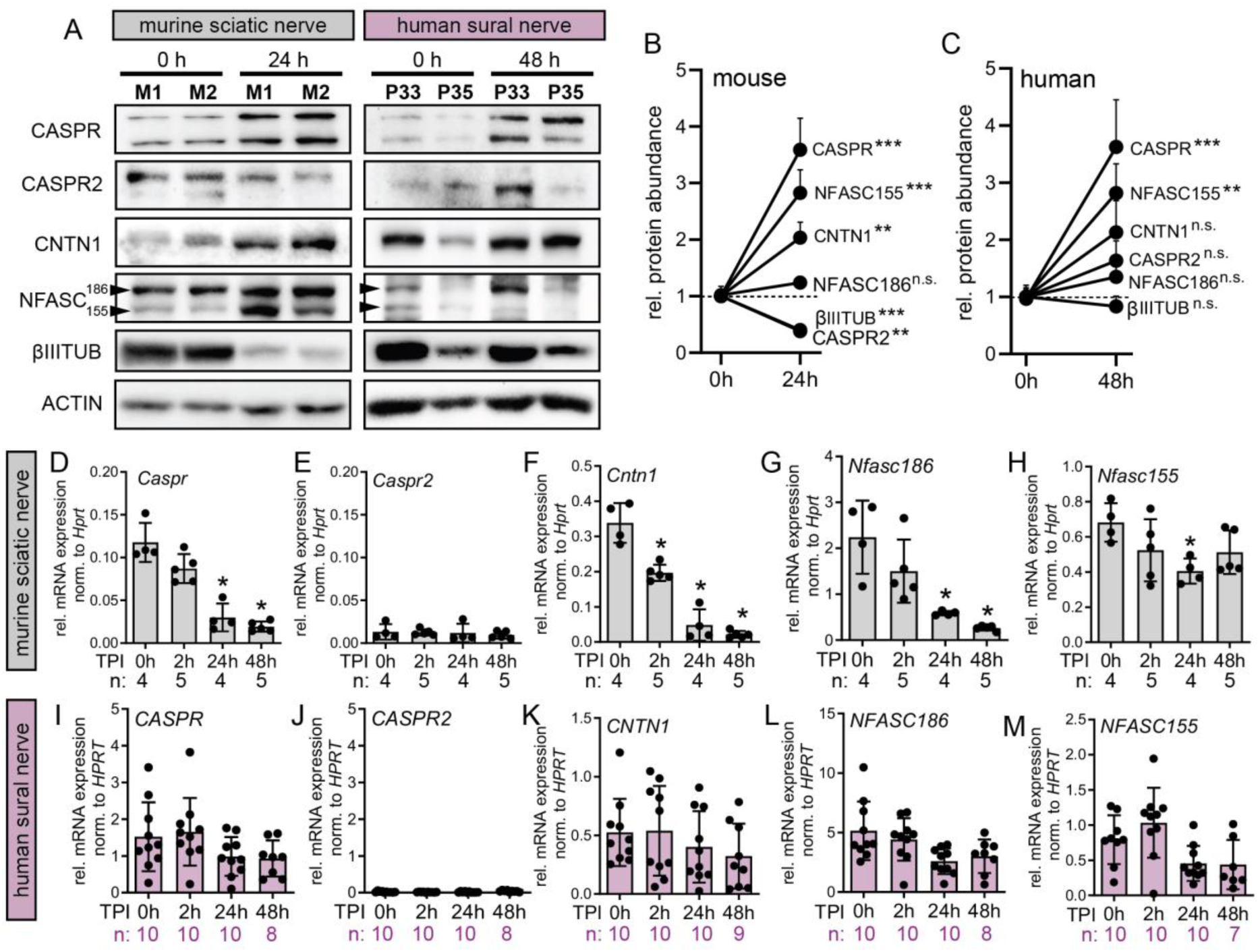
CASPR upregulation on protein level is not accompanied by transcriptional induction. (A) Immunoblots of murine sciatic and human sural nerve explants directly after dissection/surgical removal (0 h) or after *ex vivo* incubation for 24 h (murine nerves) or 48 h (human nerves). Pictures show two independent mice (M1 and M2) and two independent patient samples (Patient ID P33 and P35; see also Table 1). (B-C) Quantification of protein abundance of nodal proteins in murine and human nerves without injury (0 h) or 24h (mice)/ 48 h (human) after injury. Protein amount was calculated relative to actin abundance. The mean of all samples at 0 h was set to 1 for each protein and the corresponding mean of all samples was then calculated for each protein. (D-H) and (I-M) Gene expression analysis in murine (D-H) or human (I-M) nerve explants without injury (0h) or 2 h, 24 h and 48 h after injury and *ex vivo* incubation. Each dot in the graphs in (D-M) represents one animal or patient sample. Numbers of samples (n) are indicated below each graph. Data are presented as mean values with SD. Statistical analysis was performed using a two-sided Mann-Whitney test with *p < 0.05; **p < 0.01; ***p < 0.001. CASPR: contactin associated protein 1, CASPR2: contactin associated protein 2, CNTN1: contactin 1, NFASC: neurofascin, TPI: time post injury, βIIITUB: βIII-tublin

In addition to CASPR, we also analyzed further nodal and paranodal proteins involved in the organization of Ranvier’s nodes. This was important in order to elucidate whether upregulation is confined to CASPR alone or reflects a general regulation of axonal nodal/paranodal proteins. These proteins included the axonal proteins CASPR2, NFASC 186 and CNTN1 as well as the SC-protein NFASC 155. In murine nerves, the axonal marker βIIITUB was reduced after injury, due to axonal degradation, as was the CASPR family protein CASPR2 (Fig. 3A, B). In contrast, the axonal NFASC186 was not altered over the course of 24 h (Fig. 3A, B). Interestingly, CNTN1 and NFASC 155 which in healthy nodes build the complex that – together with CASAPR – directly mediates the axon-myelin sheath contact was also upregulated in injured murine nerves (Fig. 3A, B). Taken together, in murine nerves the CASPR/CNTN1/NFASC 155 complex seemed to be induced by injury, while other nodal proteins were unaffected or even reduced.

We then analyzed protein lysates of human nerves to be able to compare the responses. Since the previous histological results pointed at a delayed response in human compared to murine nerves (Fig. 2), we chose to use samples 48h after injury for immunoblot analysis. This confirmed the upregulation of CASPR upon injury also in human nerves (Fig. 3A, C). CNTN1 and NFASC 155 were upregulated as well, although not significantly in case of CNTN1, while other axonal proteins (CASPR2, NFASC186, βIIITUB) where not altered at this timepoint (Fig. 3A, C).

Above, upregulation of CNTN1 and NFASC 155 was noted in murine nerves and to some extent in human nerves in immunoblotting (Fig. 3A). We used histology to analyze this in more detail and confirmed a profound CNTN1 upregulation both at 24 h and 48 h after injury in both murine and human samples (Fig. S2). In addition, we found a significant reduction of βIIITUB^+^ axons over time in murine nerves, which was in line with the western blot results (Fig. S2B, D, F, M). In accordance with the generally delayed injury response in human nerves, axonal degradation was also visible in human samples, yet not significant until 48 h upon injury (Fig. S2H, J, L, N).

Next, we analyzed whether CASPR, CNTN1 and NFASC 155 protein upregulation is based on changes on transcript level. Therefore, we used murine nerve explants and analyzed the mRNA abundance of nodal/paranodal axonal proteins over time following injury. In contrast to protein levels, mRNA levels of *Caspr*, *Cntn1*, *Nfasc155* and *Nfasc186* were reduced over time (although only transiently for *Nfasc155*) compared to uninjured nerves, while *Caspr2* mRNA could not be detected in the explants at all (Fig. 3D-H). Similar results were obtained using human nerve explants, although the effects were weaker most likely due to the delayed injury response in human compared to murine nerve samples (Fig. 3I-M).

Hence, protein upregulation of CASPR and CNTN1 was not accompanied by mRNA regulation, neither in murine, nor in human nerves.

### CASPR upregulation upon injury is reduced by inhibition of protein synthesis

So far, our experiments showed that paranodal proteins CASPR and CNTN1 were upregulated on protein level in injured murine and human nerves (Figs. 1-3). Further results excluded transcriptional up-regulation contributing to this process (Fig. 3). Therefore, CASPR protein induction might be exerted by enhanced protein translation. In order to test this hypothesis, we first confirmed activation of protein biosynthesis in the *ex vivo* nerve injury model (Fig. 4). Therefore, protein lysates from murine nerves were analyzed for activated ERK (P-ERK) as well as activated S6 (P-S6_Ser235/236_ and P-S6_Ser240/244_) at different timepoints after injury (Fig. 4). Indeed, ERK was activated as early as 2 h after injury and remained active until 24 h post injury (Fig. 4A, B). Additionally, transient S6 phosphorylation – indicative of enhanced protein translation – was noted at both phosphorylation sites (P-S6_Ser235/236_ and P-S6_Ser240/244_) most prominently also at 2 h after injury (Fig. 4A, B).

**Figure 4:**
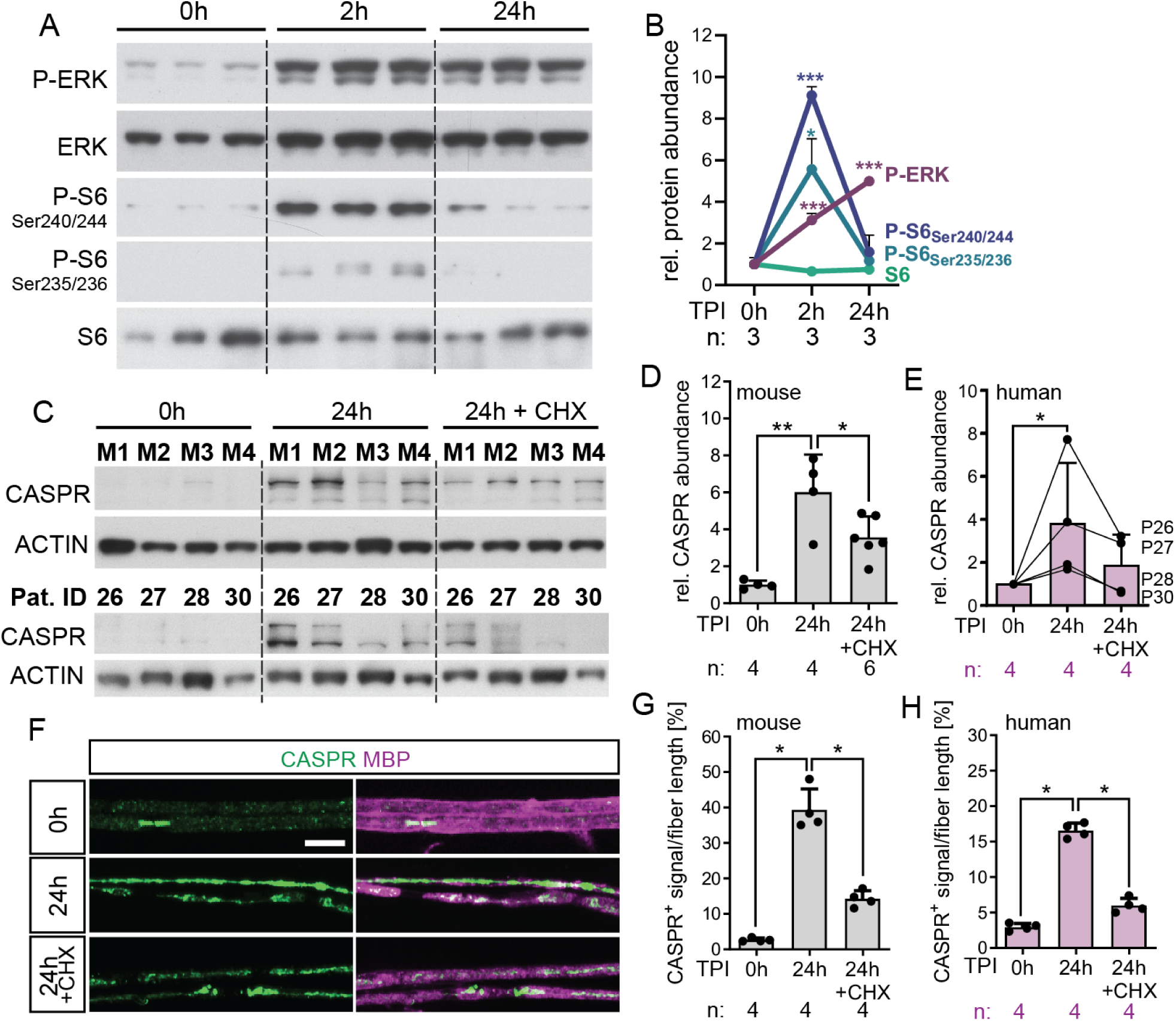
CASPR overexpression is reduced by the translation inhibitor CHX in injured axons. (A) Immunoblots of murine sciatic nerve explants uninjured (0 h) or after an *ex vivo* incubation for 2 h or 24 h. (B) Quantification of the immunoblot in (A). Protein amount was first calculated relative to ERK abundance. The mean of three biological replicates at 0 h was set to 1 for each protein and the corresponding mean of all samples was then calculated for each protein. (C) Immunoblots of murine sciatic nerve explants (four independent mice M1-M4 are depicted) and human sural nerve explants (four patient samples, patient IDs P26, P27, P28, P30) without injury (0 h) or 24 h after injury with control or CHX treatment. (D-E) Quantification of immunoblots of murine sciatic nerve (D) and human sural nerve (E) explants without injury (0 h) or 24 h after injury with control or CHX treatment. Protein amount was calculated relative to actin abundance. Connected dots in (E) indicate samples of the same patient with the patient ID depicted on the right side. (F) Teased fibers of murine sciatic nerves without injury (0 h) or 24 h after injury with control or CHX treatment. Fibers are stained for CASPR and MBP. (G-H) Quantification of CASPR^+^ signal per fiber length in teased fibers of murine sciatic nerve explants and human sural nerve explants without injury (0 h) or 24 h after injury with control or CHX treatment. Each dot in the graphs in (D-E, G-H) represents one animal or patient sample. Numbers of samples (n) are indicated below each graph. Data in (B) are presented as mean values with SEM, in (D-E, G-H) as mean with SD. Statistical analysis was performed using a two-sided t-test in (B) and two-sided Mann-Whitney test in (D-E, G-H) with *p < 0.05; **p < 0.01; ***p < 0.001. Scale bar in (F) corresponds to 10 µm. CASPR: contactin associated protein 1, CHX: cycloheximide, ERK: extracellular signal-regulated kinase, MBP: myelin basic protein, TPI: time post injury, βIIITUB: βIII-tubulin

Since protein biosynthesis was activated after nerve injury, we then proceeded to investigate, whether the observed CASPR upregulation is protein synthesis dependent. For this, the translation inhibitor cycloheximide (CHX) was added during the 24 h incubation time in order to block protein translation. Control treated nerves incubated for 24 h with DMSO showed CASPR upregulation (Fig. 4C, D). In contrast, for CHX treated nerves this upregulation was blunted (Fig. 4C, D). Finally, we used the same experimental setup and prepared teased fibers in order to confirm the observations histologically. As before, induction and re-distribution of CASPR in injured fibers was observed, which was reduced in CHX treated injured nerves (Fig. 4F-G). Of note, CHX also blocked CASPR translation in injured human nerves as shown in protein lysates and teased nerve fibers (Fig. 4C, E, H).

In sum, we showed that protein translation is induced in *ex vivo* injured nerves and local translation in axons contributes to CASPR upregulation in injured murine and human nerves.

### CASPR overexpression hinders neurite outgrowth *in vitro*

Our results above showed CASPR upregulation and redistribution in axons undergoing Wallerian degeneration. CASPR has traditionally been linked to physiological nerve impulse propagation, prompting us to explore its role in nerve pathology. Nevertheless, literature also suggests that CASPR may have inhibitory effects on neurite outgrowth on CNS neurons (16, 17).

In order to analyze how CASPR overexpression might change neuronal functions, we used primary neuronal cell cultures overexpressing either GFP or CASRP-GFP. First, we used hippocampal neurons to corroborate the published findings (16, 17) in our experimental setup. There we found that CASPR overexpression almost completely blocked neurite outgrowth (Fig. 5A-D, I-J; arrowheads indicate GFP/CASPR-GFP positive neurons). Interestingly, neurons that had not taken up the *Caspr-Gfp* expression vector during electroporation were not affected (Fig. 5C-D, arrows), indicating a cell intrinsic influence of CASPR in neurite outgrowth.

**Figure 5:**
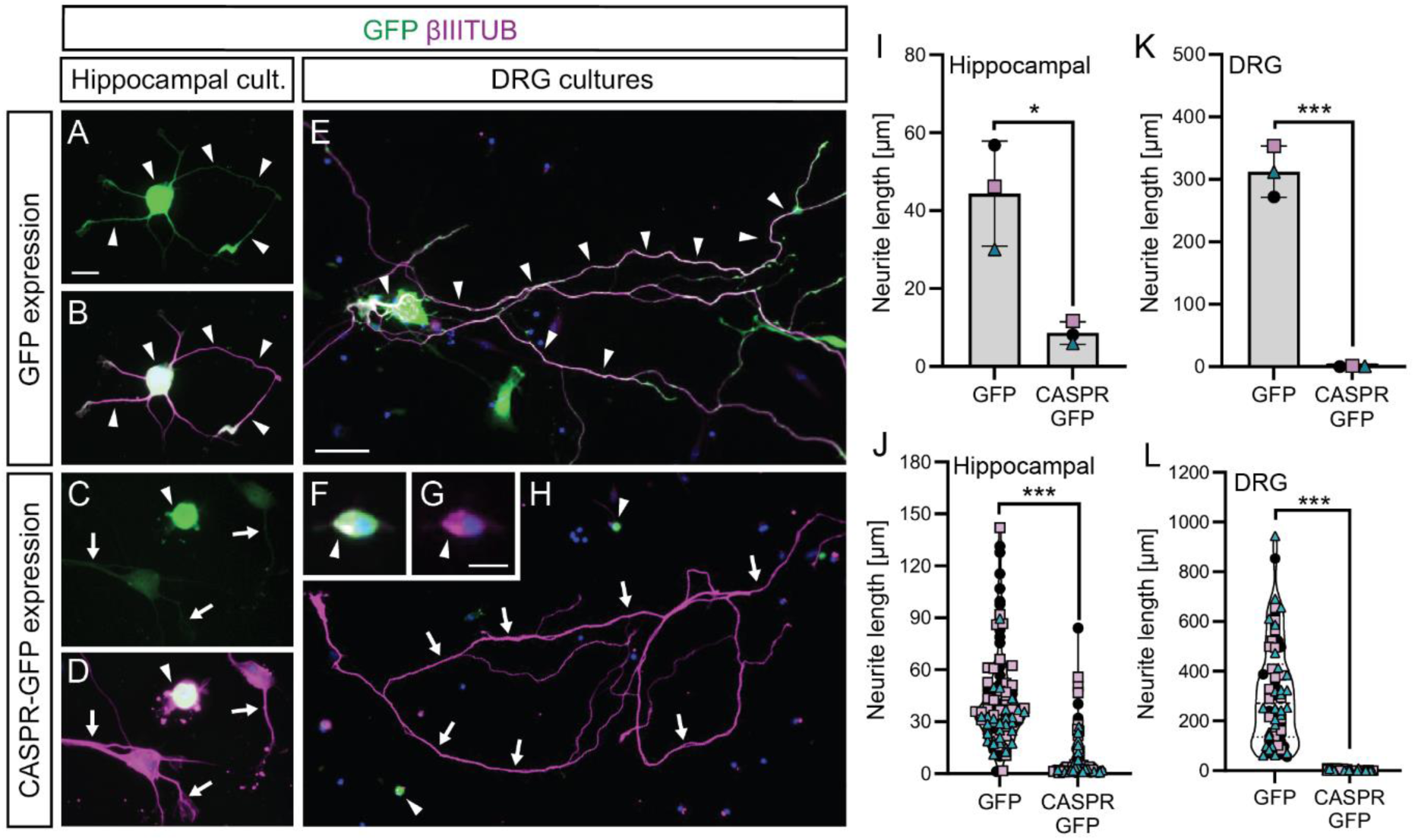
CASPR overexpression inhibits neurite outgrowth in CNS and PNS neurons. (A-D) Primary hippocampal neurons overexpressing GFP (A-B) or CASPR-GFP (C-D) stained for GFP and βIIITUB. (E-H) Primary DRG neurons overexpression GFP (E) or CASPR-GFP (F-H) stained for GFP and βIIITUB. Arrowheads point at neurons expressing GFP or CASPR/GFP while arrows point at neurons, which have not taken up the expression vectors. (I-L) Quantification of neurite lengths of hippocampal (I-J) or DRG (K-L) neurons. (I, K) Mean length of the longest neurite ± SD of three independent biological replicates, where minimum 30 neurons where quantified per condition and replicate. (J, L) Show individual neurons quantified in the three biological replicates with different symbols corresponding to the different biological replicates. Statistical analysis was performed using a two-sided t-test with *p < 0.05; **p < 0.01; ***p < 0.001. Scale bar in (A) applies for (A-D) and corresponds to 10 µm. Scale bar in (E) applies for (E, H) and corresponds to 50 µm. Scale bar in (G) applies for (F-G) and corresponds to 10 µm. CASPR: contactin associated protein 1, DRG: dorsal root ganglia, GFP: green fluorescent protein, βIIITUB: βIII-tubulin

Next, we used DRG (dorsal root ganglia) neurons to investigate if CASPR can elicit similar neurite outgrowth inhibitory effects on PNS neurons. Indeed, CASPR overexpression also blunted neurite outgrowth in DRG neurons (Fig. 5E-H, K-L; arrowheads indicate GFP/CASPR-GFP positive neurons), while neighboring CASPR-GFP negative neurons still showed normal outgrowth (Fig. 5H).

Taken together, our results show that CASPR upregulation within CNS and PNS neurons is associated with a cell intrinsic growth inhibitory signal, which limits neurite outgrowth.

### Phagocytosis of axonal debris is mediated in a CASPR depending mechanism

Thus far, we have shown that injured axons upregulate CASPR, which likely restricts axonal outgrowth in a cell-intrinsic manner. Since CASPR expression was confined to axonal segments undergoing Wallerian degeneration in our histological analysis, we next investigated whether it could also mediate a cross-talk with SCs and Mphs, therefore contributing to the overall injury response in nerves.

In nerves explants 24 h post injury, CASPR signals were frequently detected within S100β^+^ SCs (Fig. 6B-C), whereas in uninjured nerves, axonal CASPR was expectedly hardly ever observed in SCs (Fig. 6A, C). Notably, following axonal injury, degenerating axons are typically cleared through SC- and Mph-mediated phagocytosis. These observations led us to hypothesize that CASPR may function as a recognition – so-called “eat-me” – signal for SCs and Mphs in injured nerves, thereby facilitating debris clearance.

**Figure 6:**
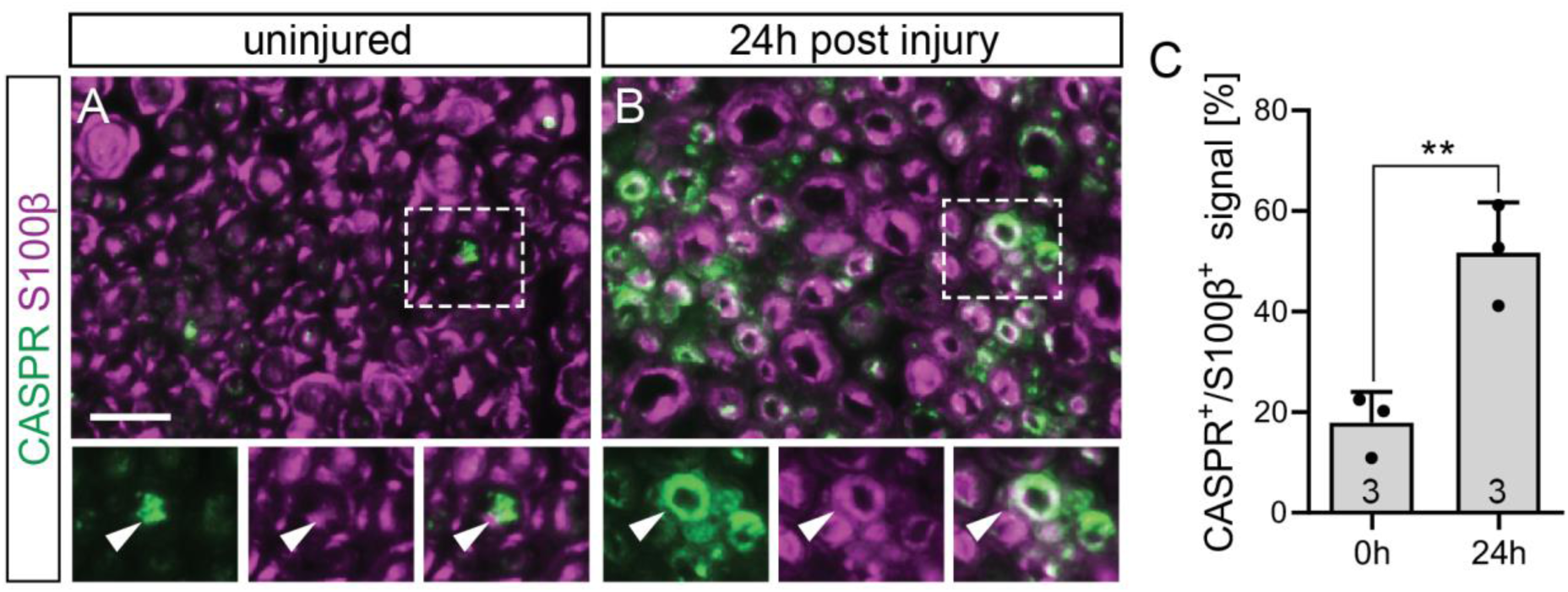
CASPR expressing axons are rapidly phagocytosed by SCs in nerves undergoing Wallerian degeneration. (A-B) Sections of murine sciatic nerve explants without injury (0 h) or after 24 h of *ex vivo* incubation. (C) Quantification of CASPR and S100β signal colocalization. Numbers of samples (n) are indicated within each bar. Data are presented as mean values with SD. Statistical analysis was performed using a two-sided t-test with *p < 0.05; **p < 0.01; ***p < 0.001. Scale bar in (A) corresponds to 10 µm. CASPR: contactin associated protein 1

To test this hypothesis, a phagocytosis assay was established using primary murine SCs and Mphs. As “feeding material” for phagocytosis, mechanically shredded nerves derived from healthy uninjured nerves were used. In addition, lysates from murine nerve explants injured for 24 h with a strong CASPR presence were employed. As a first step, we fed primary SCs of murine or human origin with this debris and evaluated the uptake of uninjured vs. injured debris. Interestingly, murine and human primary SCs phagocytosed debris originating from injured nerves more efficiently than uninjured nerve debris (Fig. 7A-D, M). To confirm that the observed debris signals within SCs originate from phagocytosis mediated uptake, we repeated the experiment with murine SCs, while treating those with LDC1267. LDC1267 is a well-established TAM receptor inhibitor, often used to block phagocytosis in different models (18, 19). Indeed, when SCs were treated with this inhibitor, nerve debris uptake was completely blocked irrespective of the type of used debris (uninjured vs. injured nerves; Fig. S3). Of note, nerve debris could not be observed within the cytoplasm of SCs due to the LDC1267 treatment, yet they were often observed right next to the SCs indicating some interaction with the cell membrane without actual uptake being possible (Fig. S3C-D, G-H, arrowheads).

**Figure 7:**
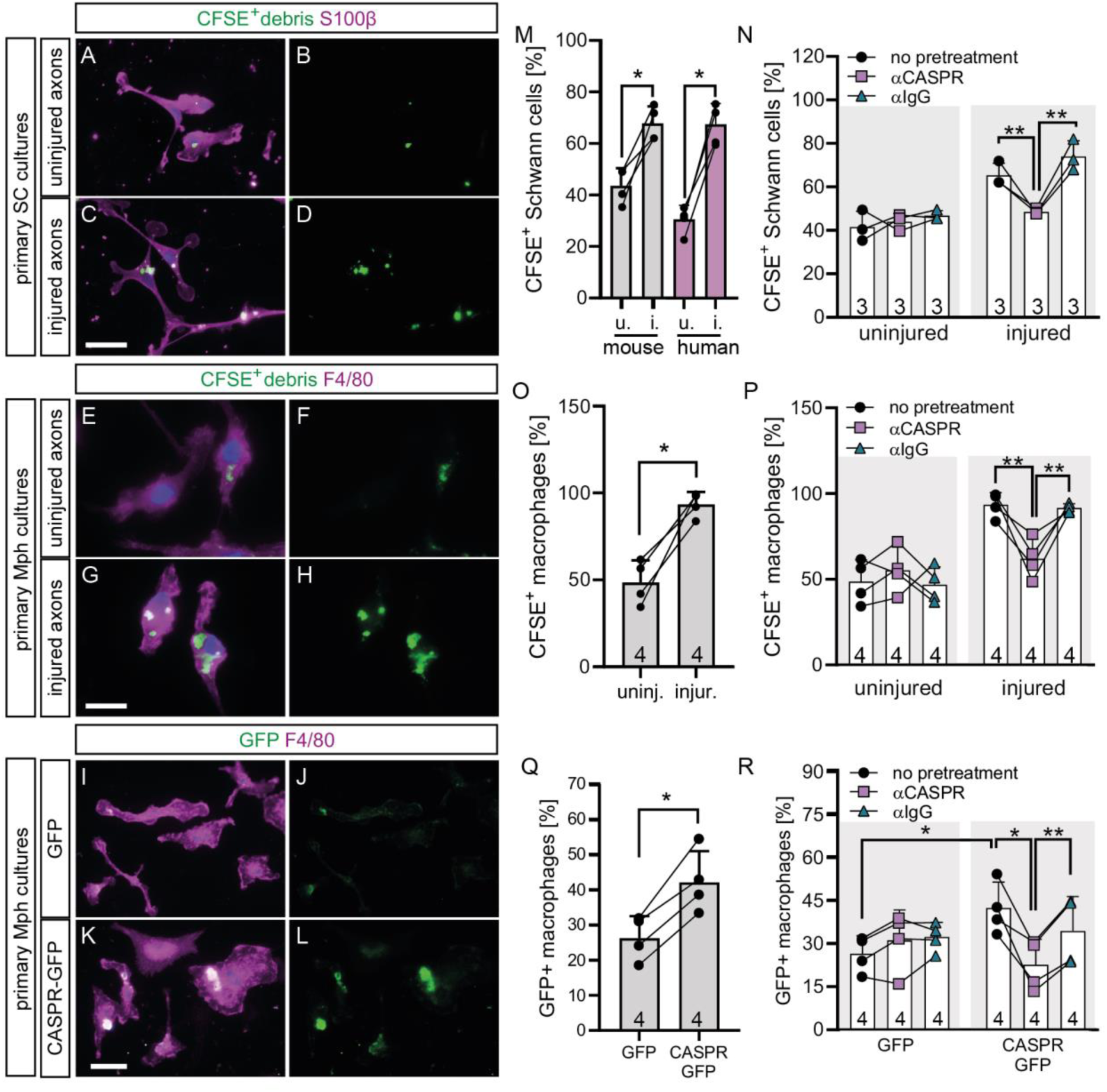
CASPR expression enhances nerve debris uptake by SCs and Mphs. (A-D) Phagocytosis assay with primary murine SC cultures. SCs were fed with CFSE labeled nerve debris from uninjured nerves (A-B) or injured nerves (C-D). (E-H) Phagocytosis assay with primary murine Mph cultures. Mphs were fed with CFSE labeled nerve debris from uninjured nerves (E-F) or injured nerves (G-H). (I-L) Phagocytosis assay with primary murine Mphs. Mphs were fed with cell debris from 3T3 cells overexpressing either GFP (I-J) or CASPR-GFP (K-L). (M, O) Quantification of the percentage of murine and human SCs (M) or murine Mphs (O) having taken up CFSE labeled nerve debris from uninjured (u./uninj.) or injured (i./injur.) nerves. (N, P) Quantification of the percentage of murine SCs (N) or murine Mphs (P) having taken up CFSE labeled nerve debris from uninjured or injured nerves, where debris was either non-treated or pre-treated with αCASPR or control αIgG antibodies. (Q) Quantification of the percentage of Mphs having taken up GFP or CASPR-GFP positive 3T3 cell debris. (R) Quantification of the percentage of murine Mphs having taken up GFP or CASPR-GFP expressing 3T3 cell debris, where debris was either non-treated or pre-treated with αCASPR or control αIgG antibodies. Numbers of samples (n) are indicated within each bar. Data are presented as mean values with SD, where each dot, square or triangle represent one biological replicate. Different conditions from one biological replicate are connected with lines. Statistical analysis was performed using a two-sided Mann-Whitney test in (M, O, P, Q, R) and two-sided t-test in (N) with *p < 0.05; **p < 0.01; ***p < 0.001. Scale bar in (C) applies for (A-D) and corresponds to 30 µm. Scale bar in (G) applies for (E-H) and corresponds to 15 µm. Scale bar in (K) applies for (I-L) and corresponds to 15 µm. CASPR: contactin associated protein 1, CFSE: carboxyfluorescein succinimidyl ester, GFP: green fluorescent protein, SC: Schwann cell, Mph: macrophage

The increased uptake of injured nerve debris compared to debris from healthy nerves suggests the presence of a signal facilitating debris uptake by SCs. To address whether such a signal might be provided by CASPR, debris from uninjured and injured nerves was pre-incubated with polyclonal anti-CASPR IgG antibodies thereby masking CASPR epitopes on the surface of axonal debris. As a control, debris was pre-incubated with non-specific IgG antibodies. Treatment of uninjured debris with anti-CASPR IgG did not alter uptake into SCs (Fig. 7N). However, treatment of debris from injured axons resulted in a significantly reduced uptake, almost to the level of uninjured debris (Fig. 7N). In contrast, control IgG did not influence debris uptake, neither from healthy, nor from injured nerves (Fig. 7N). This implies, that CASPR was most likely the mediator of enhanced debris uptake of injured nerves.

*In vivo*, SCs are the first cells at the injury site to start clearance of axonal and myelin debris (1, 4, 10). However, due to the elevated production of cytokines and chemokines by repair SCs, Mphs rapidly invade the injury site and take over debris removal (1, 4, 10). Therefore, we investigated if CASPR can also mediate debris uptake in Mphs. To test this, we repeated the above described experiment using primary murine F4/80 positive Mphs. As seen in SCs, Mphs also more readily phagocytosed debris from injured nerves as compared to debris derived from uninjured nerves (Fig. 7E-H, O). Furthermore, blocking CASPR function with anti-CASPR IgG blunted phagocytosis when used on injured nerve debris, but did not alter uptake of uninjured debris into Mphs (Fig. 7P). In contrast, use of control IgG antibodies did not have any impact on phagocytosis of uninjured or injured nerve debris. Hence, it seems that SCs and Mphs have a shared mechanism of recognizing axonal CASPR, which results in increased uptake by phagocytosing cells present in the PNS after injury.

Finally, we wanted to verify that CASPR overexpression is sufficient to induce debris uptake by phagocytes. Therefore, 3T3 cells – physiologically not expressing CASPR – were used to overexpress CASPR-GFP or GFP as a control. Subsequently, primary murine Mphs were fed with 3T3 cell debris either expressing CASPR-GFP or GFP. In line with our previous observations, debris uptake was more efficient when Mphs were fed with CASPR-GFP than GFP overexpressing 3T3 cells (Fig. 7I-L, Q). Further, treating 3T3 cell debris with anti-CASPR IgG reduced the uptake of CASPR-GFP overexpressing cells, while it did not have an influence on GFP overexpressing cells (Fig. 7R). In addition, treating 3T3 debris with control IgG antibodies did not have any influence on their uptake (Fig. 7R). Hence, CASPR overexpression seems to be sufficient to enhance phagocytosis in Mphs.

## Discussion

CASPR and its associated node proteins are essential for physiological nervous system function, including saltatory conduction and neuron-glia communication. Here, we identify a novel CASPR function in a pathophysiological context. Following peripheral nerve injury, CASPR abundance increased and CASPR redistributed along the degenerating axons, implicating a role in Wallerian degeneration (Figs. 1-3). Although this injury-induced pattern has not been previously reported in the PNS or CNS, elongated CASPR-positive domains have been observed in various neurodegenerative conditions, including demyelinating diseases, ALS, multiple sclerosis, and polyneuropathy (20–24). Our findings, along with those previous reports in brain and nerve pathologies, suggest that CASPR’s role extends beyond physiological processes to pathological conditions.

Herein, we identified two key aspects of CASPR regulation following nerve injury: (i) re-localization along axons and (ii) increased protein abundance without a corresponding rise in mRNA levels. This upregulation is mediated by translational control, as it is blocked by the ribosome inhibitor CHX (Fig. 4). Since axons are disconnected from their cell bodies in *ex vivo* nerve incubations – thereby excluding protein delivery from the cell body towards the axon – axonal *Caspr* mRNA is most likely employed by the local translation apparatus in the axons. This supports the notion that injury triggers local axonal translation of CASPR. In fact, growing evidence highlights the importance of local translation in injured nerves for effective repair and regeneration (25–27) and our findings position CASPR as a novel component of this response.

An intriguing question is what the role of CASPR in degenerating axons might be. Our results in DRG neurons (Fig. 5) together with previous studies in CNS neurons demonstrated CASPR as a negative regulator of neuronal growth (16, 17). How might this finding translate into the *in vivo* situation of nerve injury? Rapid upregulation of CASPR in injured axons probably hinders axonal outgrowth during this early phase of injury response. Although speculative at the moment, this temporal growth inhibition might be vital for the coordination of regenerative processes in injured nerves, since SC-reprogramming into repair SCs and debris removal have to precede axonal outgrowth in order to generate a growth permissive environment. In this study, we also provide evidence that CASPR acts a signal to promote phagocytic activity in SCs and Mphs (Fig. 7). This suggests that injury-induced upregulation and redistribution of CASPR along axons plays a dual role during early injury response, mediating a growth inhibition on the one hand and serving as an “eat me” signal on the other hand. Therefore, CASPR mediated phagocytosis might also provide a mechanism to remove a growth inhibiting signal.

So far, several “eat me” signals have been identified in injured peripheral nerves, including phosphatidylserine or glycosylated proteins recognized by TAM family receptors or the mannose receptor (7–10). Our phagocytosis assays support a role for TAM receptors in SC- and macrophage-mediated debris clearance, consistent with prior reports (10). This poses the question whether CASPR could promote phagocytosis via its cognate binding partners CNTN1 (on the axonal membrane) and NFASC155 (on the SC membrane) – both of which are also upregulated after injury (Fig. 3 and S2). Alternatively, CASPR might interact with some of the reported “eat me” machinery. Notably, CASPR is highly glycosylated, including mannose residues (28), suggesting a potential interaction with the mannose receptor in SCs and Mphs. Therefore, future studies should analyze whether glycosylated CASPR directly engages this receptor to trigger phagocytic activity.

Of note, data obtained in a mouse model of nerve injury were recapitulated in an *ex vivo* model of human sural nerve injury. Both, CASPR upregulation and redistribution were also conserved in injured human nerves. This points at a conserved injury signal in mammals which might be targeted by therapeutic approaches aiming at regeneration after neuronal injury.

## Materials and Methods

### Sex as a biological variable

Throughout this study, mice and human samples deriving from both sexes were used, since no differences between sexes could be observed.

### Mouse experiments

All experiments were performed using C57BL/6J mice of both sexes. Mice were maintained in groups with free access to food and water in the animal facility (12 h light-dark cycle) of Ulm University. Sciatic nerve crush was performed under general anaesthesia with isoflurane. Mice were anaesthetised and placed on a heating pad. A small skin incision was performed on the right thigh, the fascia between the biceps and the quadriceps femoris muscles were opened by blunt dissection and the sciatic nerve was crushed 5 mm proximal to the bifurcation site into tibial, sural and common peroneal nerves using haemostatic scissors. Fascia were then pushed back in place and the skin incision was sutured. Mice were sacrificed with CO_2_ after three days and nerves were dissected for histology. The uninjured sciatic nerve of the left leg served as an internal control (uninjured / 0 h).

### Human nerve samples

Human nerves were obtained from peripheral nerve transplantation surgeries conducted at the Department of Neurosurgery, Peripheral Nerve Surgery Section, at the District Hospital of Günzburg (Ulm University, Germany). Eligible participants were 18 years or older, and all procedures were performed under general anesthesia. A total of 39 patient samples were included, with a majority being male (24 out of 39; Table 1). The average patient age at the time of surgery was 52.4 years (range: 18– 75 years). Resected nerve tissue was allocated to different experiments as outlined in Table 1. In all cases, we received samples from the sural nerve. For patients P38 and P39 we additionally received samples from the brachial plexus and radial nerve respectively at positions, where the nerves had formed neuromas and were therefore surgically removed.

### Nerve explants

Murine sciatic nerves were harvested from C57BL/6J mice. Mice were sacrificed with CO_2_ and sciatic nerves of both hind limbs were dissected by a single cut at each end of the nerve with scissors. Subsequently, each nerve was divided into two pieces of approx. 1cm length each. One piece was snap frozen immediately (0 h time point). The remaining nerve was placed in tubes containing sterile Ringer solution and was incubated at 37 °C for different post-injury time points as indicated.

For human samples, one piece of each sural nerve was frozen on dry ice as soon as possible (within 5-30 minutes after collection, depending on the surgical procedure) and served as uninjured control nerve. Remaining parts of the sural nerve were cut into 1 cm pieces with a scalpel, placed into tubes with Ringer solution and incubated at 37 °C for different post-injury time points.

For cycloheximide (CHX) treatment, CHX (dissolved in dimethyl sulfoxide, DMSO) was added into the Ringer solution at a final concentration of 100 µM for 24 h. As a control, equal amounts of DMSO were used in Ringer solution. For RNA and protein isolation, samples were frozen at −80 °C either immediately or after *ex vivo* incubation for 2, 24, or 48 h. Histological samples were fixed in 4 % formaldehyde either immediately or following incubation for 2, 24, or 48 h. Similarly, samples for “teased fiber” preparation underwent fixation following the same protocol as histological samples. Thereafter, single nerve fibers were “teased” using forceps and dried on glass slides followed by staining as described previously (29).

### Primary neuronal cultures and electroporation

C57BL/6J mice were used for all neuronal cell cultures. Hippocampal cultures derived from P0-P3 (postnatal day 0-3) mice, adult DRG neurons were harvested from 8-16 weeks old mice. Isolated neurons were electroporated using the Ingenio Electroporation Solution Kit (Mirus, MIR 50108) according to the manufacturers protocol. Expression vectors used for electroporation were the pmaxGFP (Amaxa) and the Caspr-GFP described in Bonnon et al. (30), where 0.3 µg of DNA was used to electroporate one million of cells prior to plating. After electroporation, 5 x 10^3^ neurons were plated on poly-L-lysine (100 μg/ml) and laminin (5 μg/ml) coated coverslips (13 mm) in MEM media containing supplement B27 (both Gibco), glutamine and gentamycin. After 30 h in vitro, cells were fixed with 4% paraformaldehyde (PFA) and stained as described below.

### Primary SC and macrophage cultures

Sciatic nerves of C57BL/6J mice were dissected and transferred into DMEM containing 10% FCS and 1% penicillin / streptavidin (pen/strep) for eight to ten days. Medium was changed three times/week. Subsequent collagenase/dispase digestion was performed according to Haastert et al. (31) followed by trituration. The obtained single cell suspension was kept in a six well ultra-low attachment plate (Corning Costar®, 3471) over a period of 72 hours to remove fibroblasts. SCs were then seeded on poly-L-lysine/laminin coated coverslips and kept in the above-mentioned medium.

Human SCs cultures were prepared in a similar manner to murine cultures. Nerve samples derived from patients P38 (sample 1: sural nerve, sample 2: brachial plexus; see also Table 1) and P39 (sample 1: sural nerve, sample 2: radial nerve; see also Table 1).

Macrophages (BMDMs, bone marrow derived macrophages) were prepared from C57BL/6J mice as described previously (32). Briefly, humerus, femur and tibia from adult C57BL/6J mice were dissected and the bone marrow was flushed out. Cells were cultured in DMEM containing 10 % FCS, 30 % L929-cell conditioned medium, 1 % L-glutamine, 1% sodium pyruvate and 1 % pen/strep for 7 d.

### Phagocytosis assay

Primary SCs and macrophages were prepared as described above and plated on cover slips in 24-well plates for 24 h prior to debris addition.

For nerve debris preparation, two sciatic nerves (entire length from spine to the sciatic bifurcation at the knee) were dissected, pooled and snap frozen (representing uninjured nerves) or incubated *ex vivo* for 24 h prior to freezing. Subsequently, nerves were thawed on ice and mechanically dissociated in PBS using a tungsten carbide bead and a TissueLyser II (Qiagen). Nerve lysates were centrifuged, solid debris was resuspended in 350 µl CFSE (CellTrace™ CFSE, Invitrogen) staining solution in PBS and incubated for 30 min. In case of antibody pre-treatment, CFSE/PBS solution additionally contained either anti-CASPR (rabbit, Abcam, ab34151, final conc. 2 µg/ml) or rabbit IgG isotype control (Invitrogen, 02-6102, final conc. 2 µg/ml). For each well of a 24-well plate 5 µl of nerve debris were used to feed the pre-plated SCs or Mphs for 4 h. Finally, wells were washed three times with PBS to remove remaining floating debris and cells were fixed for immunocytochemistry.

For 3T3 cell debris preparation, 3T3 cells were cultured in DMEM containing 10 % FCS and 1 % penicillin/streptavidin. For transfection, 3 x 10^4^ cells per well were plated in 24-well plates and cultured for 24 h prior to transfection. Transfection was performed using Lipofectamine^TM^ transfection reagent (Invitrogen) according to the manufacturers protocol. After 48 h, cells were trypsinized and positive cells were sorted by their green florescence using the S3^TM^ cell sorter (BioRad). Sorted cells underwent two cycles of snap freezing/thawing to generate cell debris and 4 x 10^4^ dead cells per well were finally used to feed primary Mphs for 4 h. Antibody pre-treatment of 3T3 cell debris was performed as described above for nerve debris.

### Histology and Immunocytochemistry

Murine and human nerves were embedded in paraffin after fixation with 4% PFA and 5 μm microtome sections were prepared. Cells were fixed with 4% PFA and permeabilized with 0.1% Triton^TM^ X-100 (Sigma) prior to staining.

Primary antibodies used for histology or immunocytochemistry were as follows: anti-CASPR (rabbit, 1:1000, Abcam, ab34151), anti-MBP (mouse, 1:2000, BioLegend, 836504), anti-S100β (rabbit, 1:1000, Abcam, ab52642), anti-F4/80 (rat, 1:500, Biorad, MCA497GA), anti-GFP (mouse, 1:1000, Roche, 11814460001), anti-βIII-Tubulin (rabbit, 1:5000, Covance, PRB-435P) followed by anti-mouse or anti-rabbit Alexa 488 and Alexa 546 conjugated secondary antibodies (1:500 in histology/ 1:1000 in immunocytochemistry, Thermo Fisher Scientific, A-11003/ A-11008/ A-11001/ A-11003/ A-11071/ A-11039).

### Imaging quantification

Quantification of histological and cytochemical fluorescent images was performed using the ImageJ software. For each staining, a threshold was used to set to define specific staining as compared to background.

Neurite outgrowth analysis was performed by measuring the longest neurite of each neuron using the “segmented line” function of ImageJ. At least 30 arbitrary neurons were measured per condition and biological replicate. For phagocytosis assays, the CFSE/GFP stained area colocalizing with the S100β (for SCs) or F4/80 (for Mphs) staining was quantified by the automated “analyze particles” function of ImageJ. At least 30 arbitrary SCs or Mphs were measured per condition and biological replicate and the percentage of CFSE/GFP containing cells was calculated.

For histology, at least two sections per sample were analyzed and the mean value was used for quantification. For teased fibers, CASPR^+^ signals were detected for at least 10 fibers (corresponding to all fibers within the view of field of three microscopy pictures) per biological replicate. The signal was then normalized to the length of each fiber. The mean for each biological replicate was used for quantification. For quantification of SC or macrophage cultures, 10 microscopy fields (corresponding to a total of at least 50 cells) evenly distributed across the coverslip were captured and quantified. The mean of each biological replicate was used for quantification.

### Quantitative polymerase chain reaction (qPCR)

RNA was isolated from samples using TRIzol (Qiagen) and the RNeasy kit (Qiagen) according to the manufacturers protocol. Reverse transcription was performed with 0.7 μg RNA using reverse transcriptase (Promega) and random hexamers for cDNA synthesis. qPCR was performed on a Light Cycler 96 system (Roche) with the TB Green Premix Ex Taq PCR master mix (Takara). Gene Expression was calculated in relation to RNA levels of the house keeping gene *HPRT* (hypoxanthine phosphoribosyltransferase 1) to account for potential variations in total mRNA amounts. Primers used were as follows:

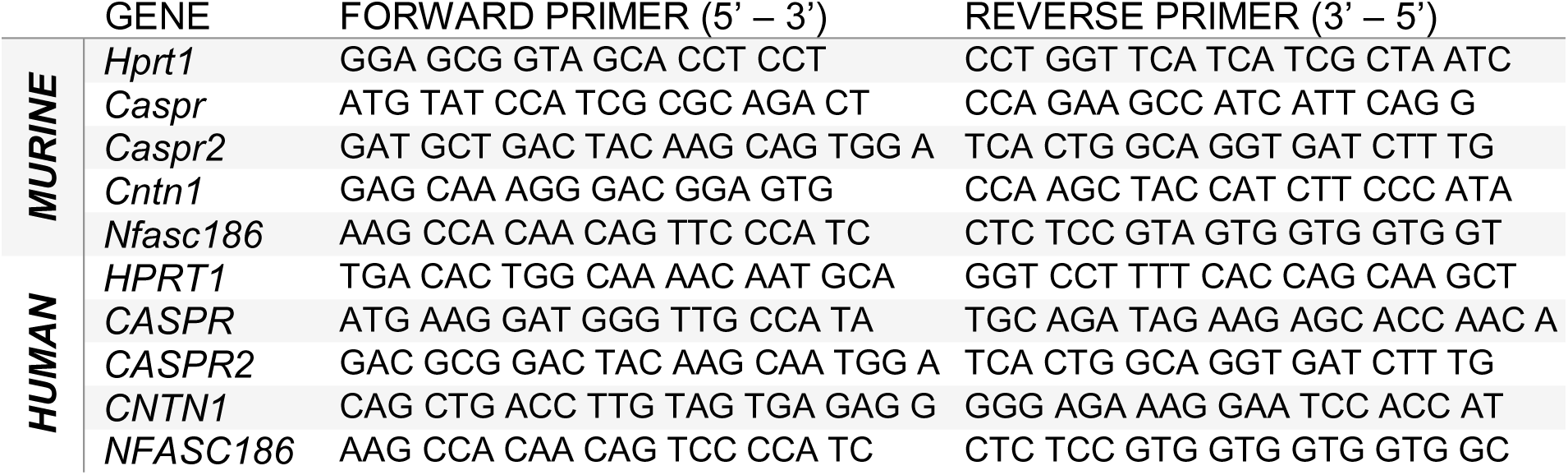

### Immunoblotting

Samples were resolved on 10% SDS–PAGE followed by transfer on PVDF membranes (Millipore). Membranes were blocked for 1 h followed by primary antibody application overnight at 4 °C: anti-CASPR (rabbit, 1:1000, Abcam, ab34151), anti-CASPR2 (rabbit, 1:500, Abcam, ab33994), anti-CNTN1 (goat, 1:500, R&D, AF904), anti-NFASC (chicken, 1:500, R&D, af3235), anti-P-ERK (rabbit, 1:1000, Cell Signaling, #4370), anti-ERK (rabbit, 1:1000, Cell Signaling, #9102), anti-S6 (rabbit, 1:1000, Cell Signaling, #2317), anti-P-S6 Ser235/236 (rabbit, 1:1000, Cell Signaling, #4858), anti-P-S6 Ser240/244 (rabbit, 1:1000, Cell Signaling, #5364), anti-Actin-HRP (mouse, 1:10.000, Santa Cruz, sc-47778). Detection of first antibodies involved horseradish-peroxidase conjugated secondary antibodies (1:3000, Cell signaling, anti-rabbit # 7074, anti-mouse # 7076, 1:2000, Millipore, anti-chicken 12-341) and the Clarity Max^TM^ Western ECL Substrate (BioRad). Membranes were either exposed on X-ray films (Fujifilm) in a dark room and developed in an Agfa film processor (X-ray developer) or developed using the ChemiDoc^TM^ imaging system (BioRad). Quantification was performed using the function “Analyze Gels” of the ImageJ software and protein abundance as normalized to actin or ERK abundance in each sample, as indicated in figures or figure legends.

### Statistical analysis

At least three biological replicates were analyzed for all experiments. Numbers (n) are indicated in each figure or figure legend. Statistical analysis of data was done with GraphPad Prism software (GraphPad Software, Inc.). Gaussian distribution was tested using the D’Agostino-Pearson omnibus normality test. Since some of the groups were not normally distributed, the non-parametric Mann-Whitney test (two-sided) was chosen if not mentioned otherwise in the figure legend. Statistical significance is provided as *, **, *** quoting P ≤ 0.05, 0.01 and 0.001, respectively. SD is provided if not mentioned otherwise.

### Study approval

All animal experiments were in accordance with institutional guidelines and German animal protection laws and were approved by the regional government authority (licence number 1515; Regierungspräsidium Tübingen, Germany).

The study for the use of human samples received approval from the local ethical review board (Nr. 91/16, 261/20, and 224/22). In accordance with the study protocol, all patients provided written informed consent before surgery.

## Author contributions

SMzR designed, conducted, analyzed and interpreted experiments and co-wrote the manuscript, A-LK conducted and analyzed experiments, AS conducted and analyzed experiments, JH conducted experiments, CK conducted experiments, DS conducted experiments, SD recruited patients for the study and prepared human samples, MP recruited patients for the study and prepared human samples, BK interpreted experiments and co-wrote the manuscript. All authors reviewed the manuscript.

## Acknowledgments

This work by SMzR and BK was supported by German Research Foundation (DFG) Project ID 251293561-SFB 1149. SMzR is additionally supported by the Medical Scientist Program of the Ulm University and the Elite PostDoc Program of the Baden-Württemberg Foundation.

## References

1. K. R. Jessen, R. Mirsky, The repair Schwann cell and its function in regenerating nerves. J Physiol 594, 3521–3531 (2016).

2. P. J. Arthur-Farraj, et al., c-Jun Reprograms Schwann Cells of Injured Nerves to Generate a Repair Cell Essential for Regeneration. Neuron 75, 633–647 (2012).

3. J. A. Gomez-Sanchez, et al., Schwann cell autophagy, myelinophagy, initiates myelin clearance from injured nerves. J Cell Biol 210, 153–168 (2015).

4. K. Hirata, M. Kawabuchi, Myelin phagocytosis by macrophages and nonmacrophages during Wallerian degeneration. Microscopy Research and Technique 57, 541–547 (2002).

5. F. Yu, et al., Phagocytic microglia and macrophages in brain injury and repair. CNS Neuroscience & Therapeutics 28, 1279–1293 (2022).

6. J. Westman, S. Grinstein, P. E. Marques, Phagocytosis of Necrotic Debris at Sites of Injury and Inflammation. Front. Immunol. 10 (2020).

7. L. Nazareth, J. St John, M. Murtaza, J. Ekberg, Phagocytosis by Peripheral Glia: Importance for Nervous System Functions and Implications in Injury and Disease. Front Cell Dev Biol 9, 660259 (2021).

8. W. Baetas-da-Cruz, et al., Schwann cells express the macrophage mannose receptor and MHC class II. Do they have a role in antigen presentation? Journal of the Peripheral Nervous System 14, 84–92 (2009).

9. V. Shacham-Silverberg, et al., Phosphatidylserine is a marker for axonal debris engulfment but its exposure can be decoupled from degeneration. Cell Death Dis 9, 1–15 (2018).

10. A. Brosius Lutz, et al., Schwann cells use TAM receptor-mediated phagocytosis in addition to autophagy to clear myelin in a mouse model of nerve injury. Proceedings of the National Academy of Sciences 114, E8072–E8080 (2017).

11. S. Einheber, et al., The axonal membrane protein Caspr, a homologue of neurexin IV, is a component of the septate-like paranodal junctions that assemble during myelination. J Cell Biol 139, 1495–1506 (1997).

12. J. C. Rios, et al., Contactin-Associated Protein (Caspr) and Contactin Form a Complex That Is Targeted to the Paranodal Junctions during Myelination. J. Neurosci. 20, 8354–8364 (2000).

13. A. Gordon, et al., Caspr and Caspr2 Are Required for Both Radial and Longitudinal Organization of Myelinated Axons. J Neurosci 34, 14820–14826 (2014).

14. S. Meyer zu Reckendorf, et al., Lipid metabolism adaptations are reduced in human compared to murine Schwann cells following injury. Nature Communications 11, 2123 (2020).

15. S. Deininger, et al., Nerve injury converts Schwann cells in a long-term repair-like state in human neuroma tissue. Exp Neurol 382, 114981 (2024).

16. V. Devanathan, et al., Cellular Form of Prion Protein Inhibits Reelin-Mediated Shedding of Caspr from the Neuronal Cell Surface to Potentiate Caspr-Mediated Inhibition of Neurite Outgrowth. J. Neurosci. 30, 9292–9305 (2010).

17. G. Suresh, R. Ramachandran, S. Sharma, K. F. Winklhofer, V. Devanathan, Novel function of Contactin associated protein 1 (Caspr 1)/ Paranodin in embryonic cortical neurons: hypoxia modulated neurite development. [Preprint] (2025). Available at: https://www.biorxiv.org/content/10.1101/2025.03.02.641114v1 [Accessed 22 April 2025].

18. Z. Zou, et al., Tyrosine Kinase Receptors Axl and MerTK Mediate the Beneficial Effect of Electroacupuncture in a Cuprizone-Induced Demyelinating Model. Evidence-Based Complementary and Alternative Medicine 2020, 3205176 (2020).

19. A. M. Gardner, et al., TAM receptor signaling dictates lesion location and clinical phenotype during experimental autoimmune encephalomyelitis. Journal of Neuroimmunology 375 (2023).

20. R. Maglemose, et al., Potassium channel abnormalities are consistent with early axon degeneration of motor axons in the G127X SOD1 mouse model of amyotrophic lateral sclerosis. Experimental Neurology 292, 154–167 (2017).

21. I. Coman, et al., Nodal, paranodal and juxtaparanodal axonal proteins during demyelination and remyelination in multiple sclerosis. Brain 129, 3186–3195 (2006).

22. A. Stojic, J. Bojcevski, S. K. Williams, R. Diem, R. Fairless, Early Nodal and Paranodal Disruption in Autoimmune Optic Neuritis. Journal of Neuropathology & Experimental Neurology 77, 361–373 (2018).

23. G. Wolswijk, R. Balesar, Changes in the expression and localization of the paranodal protein Caspr on axons in chronic multiple sclerosis. Brain 126, 1638–1649 (2003).

24. L. Appeltshauser, et al., Super-resolution imaging pinpoints the periodic ultrastructure at the human node of Ranvier and its disruption in patients with polyneuropathy. Neurobiology of Disease 182, 106139 (2023).

25. A. Vaquié, et al., Injured Axons Instruct Schwann Cells to Build Constricting Actin Spheres to Accelerate Axonal Disintegration. Cell Reports 27, 3152–3166.e7 (2019).

26. S. Koley, M. Rozenbaum, M. Fainzilber, M. Terenzio, Translating regeneration: Local protein synthesis in the neuronal injury response. Neurosci Res 139, 26– 36 (2019).

27. A. Pacheco, T. T. Merianda, J. L. Twiss, G. Gallo, Mechanism and role of the intra-axonal Calreticulin translation in response to axonal injury. Experimental Neurology 323, 113072 (2020).

28. L. Gollan, D. Salomon, J. L. Salzer, E. Peles, Caspr regulates the processing of contactin and inhibits its binding to neurofascin. J Cell Biol 163, 1213–1218 (2003).

29. J. Wen, L. Li, D. Tan, J. Guo, Preparation of Teased Nerve Fibers from Rat Sciatic Nerve. Bio Protoc 7, e2572 (2017).

30. C. Bonnon, L. Goutebroze, N. Denisenko-Nehrbass, J.-A. Girault, C. Faivre-Sarrailh, The Paranodal Complex of F3/Contactin and Caspr/Paranodin Traffics to the Cell Surface via a Non-conventional Pathway*. Journal of Biological Chemistry 278, 48339–48347 (2003).

31. K. Haastert, C. Mauritz, S. Chaturvedi, C. Grothe, Human and rat adult Schwann cell cultures: fast and efficient enrichment and highly effective non-viral transfection protocol. Nature Protocols 2, 99–104 (2007).

32. P. Pfänder, A.-K. Eiers, U. Burret, S. Vettorazzi, Deletion of Cdk5 in Macrophages Ameliorates Anti-Inflammatory Response during Endotoxemia through Induction of C-Maf and Il-10. International Journal of Molecular Sciences 22, 9648 (2021).

