## Supplementary Information for "CASPR facilitates clearance of degenerating axons by Schwann cells and macrophages after peripheral nerve injury"

[1] Institute of Neurobiochemistry
Ulm University
Albert-Einstein-Allee 11
89081 Ulm
Germany

[2] Peripheral Nerve Surgery Unit
Department of Neurosurgery
Ulm University
District Hospital
89312 Günzburg
Germany

*Co-Corresponding authors:
Sofia Meyer zu Reckendorf and Bernd Knöll



The authors have declared that no conflict of interest exists.

### Supplementary Materials and Methods

*Histology*

Additional primary antibody used in supplementary figure S2: CNTN1 (goat, 1:500, R&D Systems, AF904).

*LDC1267 treatment of primary murine SCs*

Preparation of primary murine SCs and phagocytosis assay was performed as described in the main manuscript. LDC1267 (Sigma-Aldrich) was dissolved in DMSO and added to the cell culture media at a final concentration of 0.1 µM or 1 µM for 30 min prior to adding the nerve debris. Phagocytosis assay was the continued for 4 h as described in the main manuscript.

*Study approval*

All animal experiments were in accordance with institutional guidelines and German animal protection laws and were approved by the regional government authority (licence number 1515 for sciatic nerve lesion, licence number 1389 for facial nerve lesion; Regierungspräsidium Tübingen, Germany).

### Supplementary Figures

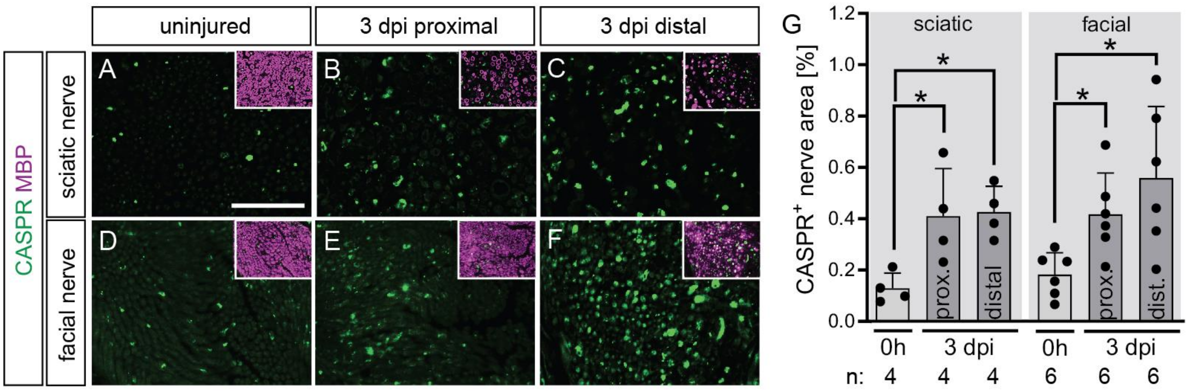


**Figure S1: CASPR upregulation upon injury is conserved in different peripheral nerves**

(A-C) Cross sections of murine sciatic nerves without injury (A) or 3 days post injury proximal (B) or distal (C) to the injury site. (D-F) Cross sections of murine facial nerves without injury (D) or 3 days post injury proximal (E) or distal (F) to the injury site. Pictures show CASPR staining while inserts depict the overlay of CASPR with MBP. (G) Quantification of the CASPR positive nerve area in uninjured vs. injured nerves at a position proximal or distal to the injury at 3 days post injury. Each dot in the graph represents one animal. Numbers of samples (n) are indicated below the graph. Data are presented as mean values with SD. Statistical analysis was performed using a two-sided Mann-Whitney test with *$p < 0.05$; **$p < 0.01$; ***$p < 0.001$. Scale bar in (A) applies to (A-F) and corresponds to 50 µm.

CASPR: contactin associated protein 1, dpi: days post injury

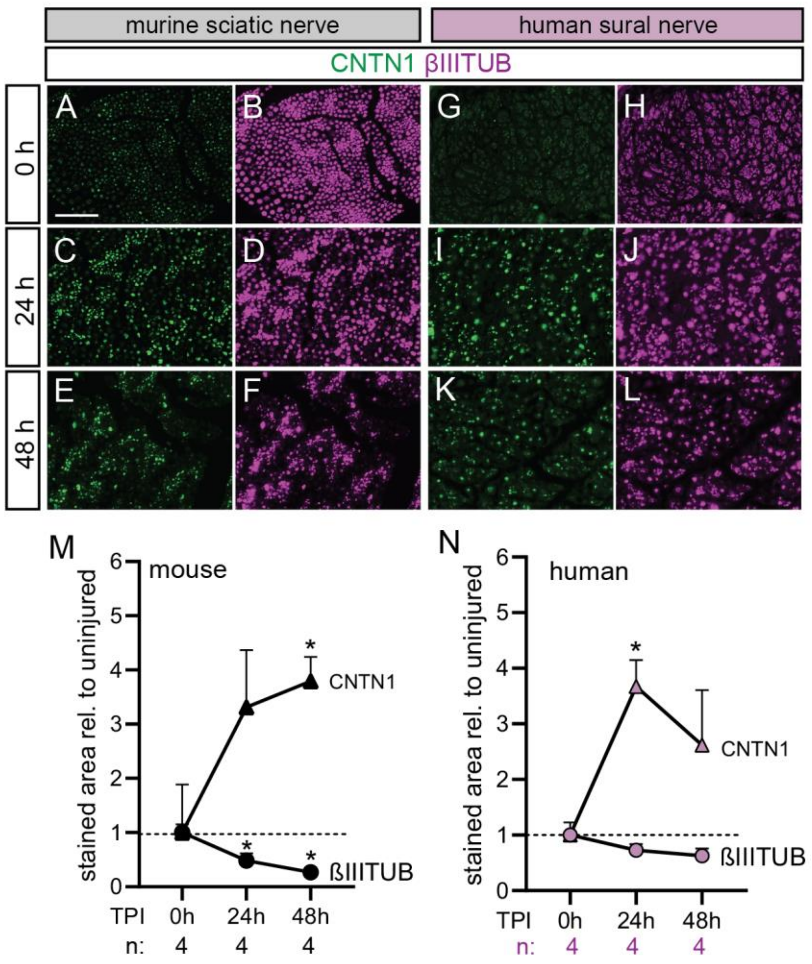


**Figure S2: CNTN1 is upregulated together with CASPR in murine and human nerve explants upon injury**

(A-L) Cross sections of murine sciatic (A-F) and human sural (G-L) nerve explants without injury (0 h) or after an ex vivo incubation for 24 h or 48 h. Pictures show CNTN1 (A, C, E, G, I, K) and βIIITUB (B, D, F, H, J, L) staining. (M, N) Quantification of the CNTN1 or βIIITUB stained area for each time point after injury, normalized to the area in the uninjured nerve in murine (M) and human (N) nerves. Numbers of samples (n) are indicated below the graph. Data are presented as mean values with SD. Statistical analysis was performed using a two-sided Mann-Whitney test with $*p < 0.05$; $**p < 0.01$; $***p < 0.001$. Stars indicate significance between the given time point compared to the 0 h time point. Scale bar in (A) applies to (A-L) and corresponds to 50 µm.

CASPR: contactin associated protein 1, CNTN1: contactin 1, TPI: time post injury, βIIITUB: βIII-tubulin

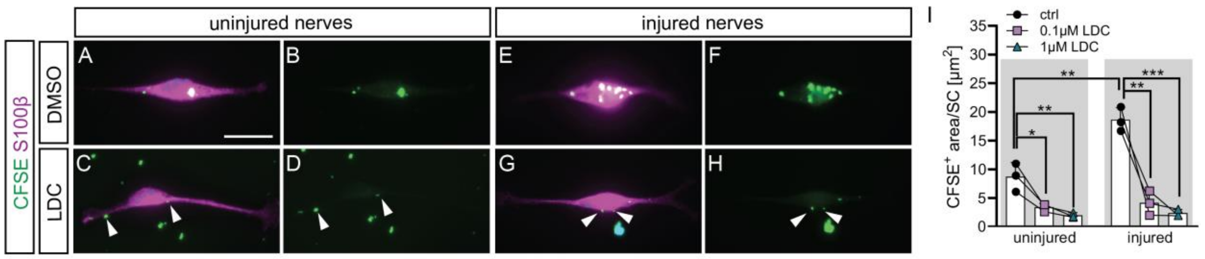


**Figure S3: Nerve debris uptake is mediated by TAM receptors in cultured SCs**

(A-H) Phagocytosis assay with primary murine SCs. SCs were fed with CFSE labeled nerve debris from uninjured (A-D) or injured nerves (E-H), while being treated with vehicle control (DMSO; A-B, E-F) or TAM receptor inhibitor LDC1267 (C-D, G-H). Arrowheads point at debris localized directly at the cell border but not within the cell. (I) Quantification of the percentage of SCs having taken up CFSE positive nerve debris under control treatment (DMSO) or treatment with two concentrations of LDC1267. Three biological replicates were analyzed. Data are presented as mean values with SD, where each dot, square or triangle represent one biological replicate. Different conditions from one biological replicate are connected with lines. Statistical analysis was performed using a two-sided t- test with *$p < 0.05$; **$p < 0.01$; ***$p < 0.001$. Scale bar in (A) applies for (A-H) and corresponds to 20 µm.

CFSE: carboxyfluorescein succinimidyl ester, DMSO: dimethyl sulfoxide, LDC: TAM receptor inhibitor LDC1267 SC: Schwann cell, TAM receptors: group of Tyro-3, AXL, MerTK receptors
